# Comparative transcriptomic analyses reveal differences in the responses of diploid and triploid eastern oysters to environmental stress during a summer mortality event

**DOI:** 10.1101/2024.01.31.578323

**Authors:** Rujuta V. Vaidya, Sarah Bodenstein, Dildorakhon Rasulova, Jerome F. La Peyre, Morgan W. Kelly

**Affiliations:** Department of Biological Sciences, Louisiana State University, Baton Rouge, LA, USA; School of Animal Sciences, Louisiana State University Agricultural Center, Baton Rouge, LA 70803, United States of America

## Abstract

Triploid oysters are commonly used as the basis for production in the aquaculture of eastern oysters along the U.S East and Gulf of Mexico coasts. While they are valued for their rapid growth, incidents of triploid mortality during summer months have been well documented in eastern oysters, especially at low salinity sites. We compared global transcriptomic responses of diploid and triploid oysters bred from the same three maternal lines and outplanted to a high (annual mean salinity = 19.4 ± 6.7) and low (annual mean salinity = 9.3 ± 5.0) salinity site at the onset of mortality event in summer of 2021 to test for effect of parental contribution on triploid performance. We compared transcriptomic responses of same diploid and triploid oysters to test for ploidy specific differences in gene expression in response to high and low salinity sites and tested for instances of aneuploidy in experimental triploid oyster lines. Maternal parentage did not affect triploid mortality, but a strong effect of hatchery conditions (cohort) was observed. We detected a higher number of DEGs in response to outplant sites (salinity) and cohorts, indicating stronger influence of these two factors on triploid mortality. At the low salinity site where triploid oysters experienced high mortality, we observed downregulation of transcripts related to calcium signaling (Calmodulin-a, Histidine-rich calcium binding protein (HRC), and cadherin EGF LAG seven-pass G-type receptor 1(CELSR1)), ciliary activity (axonemal and cytoplasmic dyneins), and cell cycle check points (CDK1, HAUS augmin-like complex subunit 3, and MAPKKK, MCM7, SMCs, RTEL1). These transcripts suggest dampening of the salinity stress response and problems during cell division as key contributors to elevated summer mortality in triploid oysters. No instances of aneuploidy were detected in our triploid oyster lines.

## 1. Introduction

Polyploidy is the heritable condition in which a typically diploid cell or organism inherits an additional set (or sets) of chromosomes (de Sousa et al., 2016). Instances of polyploidy occur when chromosomes fail to separate normally during cell division, resulting in gametes that have unequal distribution of chromosomes. Having extra sets of chromosomes affects many structural and physiological aspects of the cell – polyploid individuals are generally bigger in size, have faster growth rates, and offer better yields in commercially valuable species (Piferrer et al., 2009). The prospect of bigger marketable product has generated great interest in the generation of polyploids in agriculture and aquaculture, including in commercially valuable bivalves like the eastern oyster. Generation of triploid and tetraploid eastern oysters was first reported in 1981 (Stanley et al., 1981) and in later years, several standardized protocols for hatchery production of triploid and tetraploid oysters have been published (Wadsworth et al., 2019; H. Yang, 2022, Hudson, K., 2019) and adapted by oyster farms as their primary basis for production. Compared to the natural diploid oysters, triploid oysters generally grow faster, and have reduced gametogenesis as a consequence of their triploidy (Allen Jr. and Downing, 1986, 1990; Wadsworth et al., 2019a; Matt and Allen, 2021). Unlike diploid oysters, who invest considerable energy reserves in reproduction, triploids oysters instead channel that energy into their growth, resulting in better meat quality during summer (Barber and Mann, 1991; Degremont et al., 2012;Walton et al., 2013). All of these ‘triploid advantages’ have made them a desirable alternative to natural diploid oysters, and triploids have been adapted by many hatcheries as their primary basis for production (J. Brianik & Allam, 2023)

However, in recent years there have been reports of triploid eastern oysters experiencing higher mortality than diploid oysters during spring and summer months. This phenomenon, collectively termed as ‘triploid mortality’ has been observed along the US mid-Atlantic and Gulf of Mexico (GoM) coast.(George et al., 2023; Houssin et al., 2019; Matt et al., 2020; Wadsworth et al., 2019). Several factors like unfavorable environmental conditions (salinity and temperature), parental stocks, and disrupted gene regulation and stress response in triploid oysters have been suggested to contribute to triploid mortality, however the exact underlying cellular mechanisms causing these triploid mortality events remain unresolved (Auger et al., 2005; Callam et al., 2016; Ching et al., 2010; R. E. Hand et al., 2004; Marshall et al., 2021; Marshall & Sandra M Casas, 2021; Swam et al., 2022). Additionally, chromosome losses in polyploid organisms occur are more frequent in polyploid organisms (Comai, 2005), and instances of aneuploidy have been reported in early cell divisions of polyploid oysters (de Sousa et al., 2016; Z. Wang et al., 1999). Losing a part of or a whole chromosome can cause major disruptions in patterns of gene expression and regulation, which can affect the susceptibility of triploid oysters to unfavorable environmental conditions.

In our study, we compared transcriptomic responses of diploid and triploid oysters bred from the same three maternal lines and outplanted to a high (annual mean salinity = 19.4 ± 6.7) and low (annual mean salinity = 9.3 ± 5.0) salinity site at the onset of mortality event in summer of 2021 to test for effect of parental contribution on triploid performance. To investigate whether triploid oysters show disrupted gene regulation and stress response in response to unfavorable environmental conditions, we compared the global transcriptomic response of the same diploid and triploid oysters to high and low salinity sites. Instances of aneuploidy in triploid oysters used in the experiment were also tested using methods described in Griffiths et al., 2017 to our transcriptomic dataset. Environmental data (salinity and temperature) and morphological measurements including growth, *Perkinsus marinus* (dermo) infection prevalence, and mortality were also performed and are explained in Bodenstein et al., 2023. Interestingly, the mortality data reported in Bodenstein et al., 2023 showed no effect of maternal stocks on triploid survival, but showed strong effect of hatchery conditions (referred as cohorts), with one of the two hatcheries used for rearing oyster lines showing higher triploid mortality. We therefore decided to include the two hatchery conditions in our pairwise diploid vs triploid comparisons instead of maternal lines.

## 2. Methods

### a) Oyster crossing and outplanting

The detailed cross strategy and outplanting designs are explained in Bodenstein et al., 2023 and is summarized in Figure 1. Briefly, to establish diploid parental broodstocks adapted to different salinity regimes, oysters were collected from each of three Louisiana public oyster grounds that had different salinity regimes: Calcasieu Lake (CL, annual mean salinity = 16.2 ± 2.8 [± standard deviation (SD), n= 10, 2009–2018]), Sister Lake (SL, annual mean salinity = 11.2 ± 5.5), Vermilion Bay (VB, annual mean salinity = 7.4 ± 1.6). The tetraploids used in this study were LSU 4DGNL17 (maintained at Louisiana Sea Grant Oyster Research Farm) and AU 4MC18 (maintained at the Grand Bay Oyster Park, Alabama). Oysters were conditioned and spawned in June 2019 at Mike C. Voisin Oyster Hatchery (LSU cohort) and at the Auburn University

**Figure 1:**
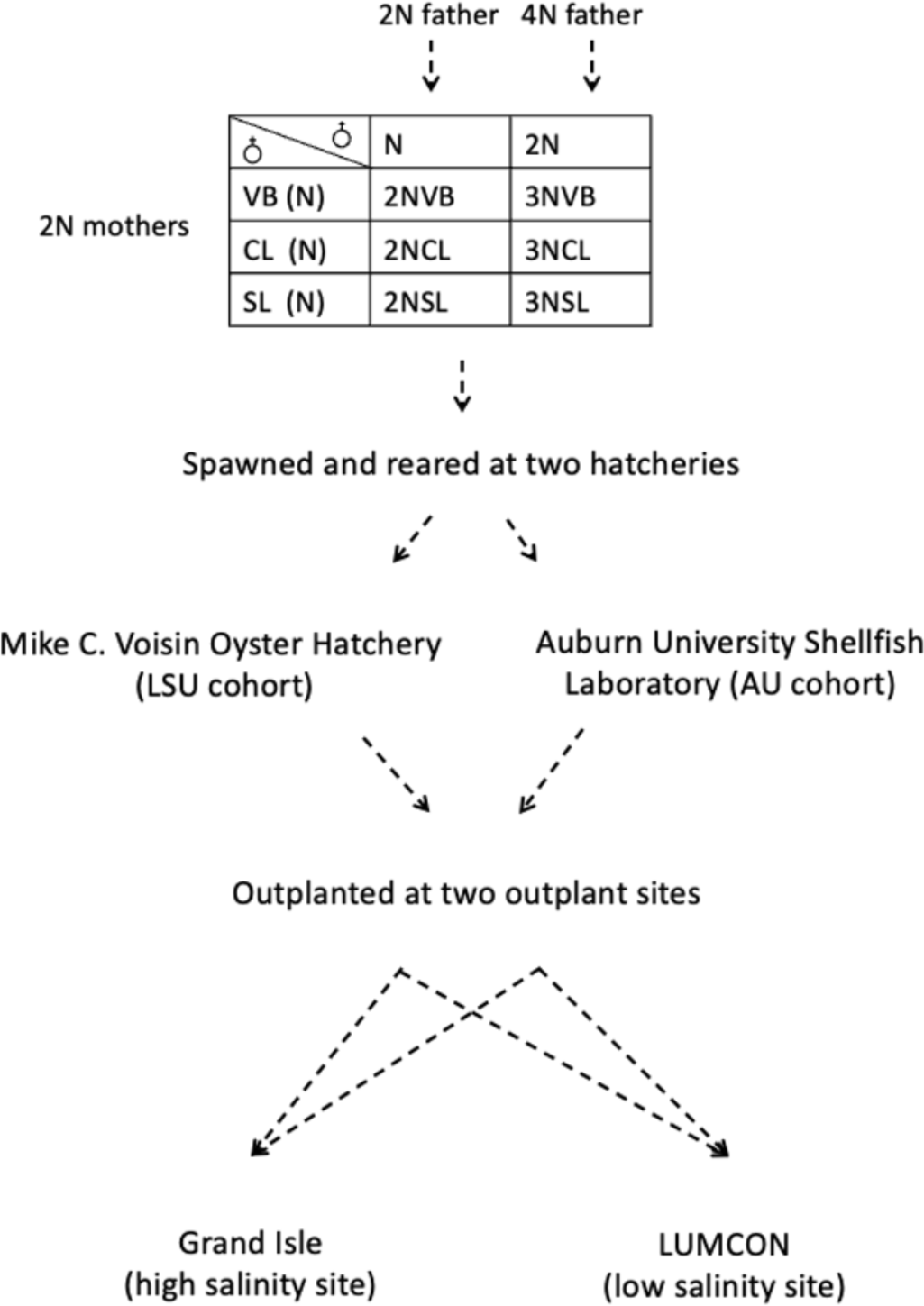
Cross strategy used to generate diploid and triploid lines for this study. Detailed cross strategy and number of oysters used for each cross are described in Bodenstein et al., 2023.

Shellfish Laboratory (AU cohort) in Dauphin Island, AL in July 2019 as described in Bodenstein et al., 2023. Depending on where they were spawned, F1 progeny was sorted into LSU (Louisiana State University) or AU (Auburn) cohorts. A total of six crosses were produced at each hatchery. The diploid F1 crosses were produced by crossing male and female oysters from each wild broodstock population (CL, SL, VB) and the triploid F1 crosses were produced by crossing females from wild broodstock populations with males from two tetraploid lines (one from each hatchery, Fig. 1). Diploid and triploid crosses were generated from the same females at the Auburn hatchery and were therefore half-siblings. However, at the LSU hatchery, diploid and triploid crosses were not half-siblings and, in certain cases, were produced on separate days. Oysters were reared and transported to their final outplant sites as described in Bodenstein et al., 2023. Oysters from both cohorts were outplanted to a low (near the Louisiana Universities Marine Consortium (LUMCON), annual mean salinity = 9.3 ± 5.0) and a moderate salinity site (Grand Isle, annual mean salinity = 19.4 ± 6.7) in Louisiana. Oysters from Auburn cohort were outplanted in November 2019 at the two sites and oysters from LSU cohort were outplanted at Grand Isle in December 2019 and at LUMCON in January 2020. At each site, four replicate baskets containing 80 oysters each were outplanted for each of the six crosses of each ploidy in the two cohorts, a total of 48 baskets per site. Growth and mortality measurements were performed monthly till November 2020 and are described in Bodenstein et al., 2023.

### b) RNA sequencing

On 8^th^ July, 2020, eight individuals per cross were sampled at both outplant sites. Approximately 0.5-cm^2^ piece of gill tissue was dissected in the field from each individual and preserved with Invitrogen RNAlater. Total RNA was extracted using an E.Z.N.A. Total RNA Kit I (Omega BIO-TEK; VWR catalog no. 101319) following the manufacturer’s instructions. The yield and quantity were assessed using a NanoDrop 2000 spectrophotometer. Total RNA extracted from the 192 individuals were sent to the University of Texas at Austin’s Genomic Sequencing and Analysis Facility where RNA quality control was confirmed using a 2100 Agilent Bioanalyzer on a Eukaryote Total RNA Nano chip and libraries were produced using the Tag-sequencing approach (Meyer et al., 2011). The resulting 192 libraries were sequenced equally across two lanes of an Illumina HiSeq 2500 platform, with 100-bp single-end reads. Adapter sequences were trimmed using trimmomatic (version 0.39) (Bolger et al., 2014) and base pairs with quality scores below 35 were removed. The trimmed reads were then mapped to the *C. virginica* reference genome (Gómez-Chiarri et al., 2015) with known haplotigs removed (https://github.com/jpuri tz/ Oyste rGenomeProject/tree/maste r/Haplo tig_Masked_Genome) using star rna-seq aligner (single pass option). Reads were mapped to gene features with the options (-- quantMode GeneCounts –outFilterScoreMinOverLread 0.50 -- outFilterMatchNminOverLread 0.50) to adjust for poly-A tail contamination (Sirovy et al., 2021) to finally generate a count matrix using ReadsPerGene.out.tab output (supplementary file 1).

### c) Differential Gene Expression (DEG)

To maximize the true positive rate and specificity, we used both parametric and non-parametric tests to obtain a single consensus list of DEGs (Costa-Silva et al., 2017) for our pairwise comparisons. DEGs were calculated using DESeq2 (v 1.24.0) (Love et al., 2014), EdgeR (Robinson et al., 2010), and NOISeq (Tarazona et al., 2011) and the R package VennDiagram was used to find consensus lists of DEGs detected by all three methods for all comparisons.

DEGs were identified using a false discovery rate (FDR)-adjusted p-value cutoff of 0.01. Since mortality data reported in Bodenstein et al., 2023 showed that maternal parentage did not have any effect on triploid mortality, we did not include dams in our pairwise DEG comparisons. The mortality data showed strong effect of cohort on triploid survival and was therefore incorporated in our DEG analysis. We performed pairwise comparisons of diploid and triploid oysters across cohorts and ploidy. Gene ontology enrichment was also performed using GO_MWU (Wright et al., 2015), using the Fisher’s Exact Test (p<0.05) the scripts available at: https://github.com/z0on/GO_MWU/blob/master/GO_MWU.R. Differentially expressed genes (FDR<0.05) were calculated using pairwise comparisons in EDGER, NOISeq, and DESeq2. DEG lists from all three approaches were compared and only genes that were detected by all three approaches were used as DEGs for downstream analysis.

### d) Weighted Gene Correlation Network Analysis (WGCNA)

The R package WGCNA (Langfelder & Horvath, 2008) was used to identify clusters of genes with highly correlated expression patterns. We filtered out genes with fewer than five counts per million in 80% of all samples to remove lowly expressed features retaining only samples with greater than five counts per million in 80% of all samples. The WGCNA was run using 9172 genes that passed this criterion, using a soft threshold value of 12, a minimum module size of 30, and a signed adjacency matrix. The network was then correlated to site, cohort, and ploidy, with variables converted to binary format. Module eigengenes, defined as first principle component of a given module (Langfelder & Horvath, 2008) were visualized using ggplot2 and GO enrichment analysis was conducted using GO_MWU. To measure the magnitude of the difference between transcriptomic responses of our diploid and triploid oysters, we weighted the samples by the percent of variation explained by each axis and then calculated the shift (Euclidean distances) in multivariate space between diploid and triploid samples matched for their cohort, site, and dams.

### e) Testing for aneuploidy in triploid oysters

We also assessed for instances of aneuploidy across all 10 oyster chromosomes. To do that, we first categorized our samples into diploid and triploid pairs by matching their site, cohort, and dam, i.e. a diploid oyster from Auburn cohort and VB dam broodstock and was outplanted to Grand Isle was paired with a triploid oyster from Auburn cohort and VB dam broodstock and was outplanted to Grand Isle. For every pair, we then plotted normalized log transformed Counts Per Millions (CPM) values such that each point on the plot represented a transcript and X and Y coordinates of every point were CPM values of that transcript in the diploid and triploid oyster respectively. For each pair compared, ten such plots (one for each chromosome) were made, and the slope of the regression line was calculated. As the CPM values were normalized to the library sizes, change in the slope for any chromosome for a particular sample pair was considered as an indicator of aneuploidy occurring at that chromosome (Griffiths et al., 2017; Stamoulis et al., 2019; Stingele et al., 2012; Zhao et al., 2017).

## 3. Results

### 3.2 Principal Component Analysis and aneuploidy analysis

Detailed mortality and growth measurements are described in Bodenstein et al.,2023. Overall, triploids showed higher mortality at low salinity site (LUMCON) (Fig. 2). At both high and low salinity sites, the cumulative mortality of triploids was higher than diploids, and triploids from LSU cohort experienced higher cumulative mortality than AU cohort. Maternal stock (dams) did not influence cumulative mortality of triploids at either site. A principal component analysis (PCA) was performed to visualize broad patterns of gene expressions. Raw counts data for diploid and triploid oysters was first normalized using variance stabilizing transformation (VST) function in DESeq2 package and PCA was performed using plotPCA function in DESeq2 package. The first two principal component axes accounted for 61% and 6% variance in samples respectively (Fig. 3a). Out of the four variables, outplant site had the greatest effect on sample clustering, followed by cohort and ploidy. No clustering was observed based on dams. The magnitude of transcriptomic response for diploid and triploid oysters was visualized using PCA as per Thomas et al., 2022. Briefly, we extracted PC1 and PC2 values for each sample (represented by a point in the PCA plot) to calculate Euclidean distances between each diploid and triploid sample pair matched for their cohort, site, and dams. The distances were plotted as separate box plots for Grand Isle and LUMCON sites and color coded by their cohorts. Overall, the difference in magnitude was higher in diploid vs triploid oyster comparisons at LUMCON site, regardless of their cohorts. To check if the differences in mortality were a result of aneuploidy in triploid oysters, we plotted normalized log transformed CPM values of diploid and triploid oysters on X and Y axis respectively. Slope values for all ten chromosomes for each sample pair are summarized in supplementary table 1 . We do not see change in slopes of any chromosomes for any of our sample pairs.

**Figure 2:**
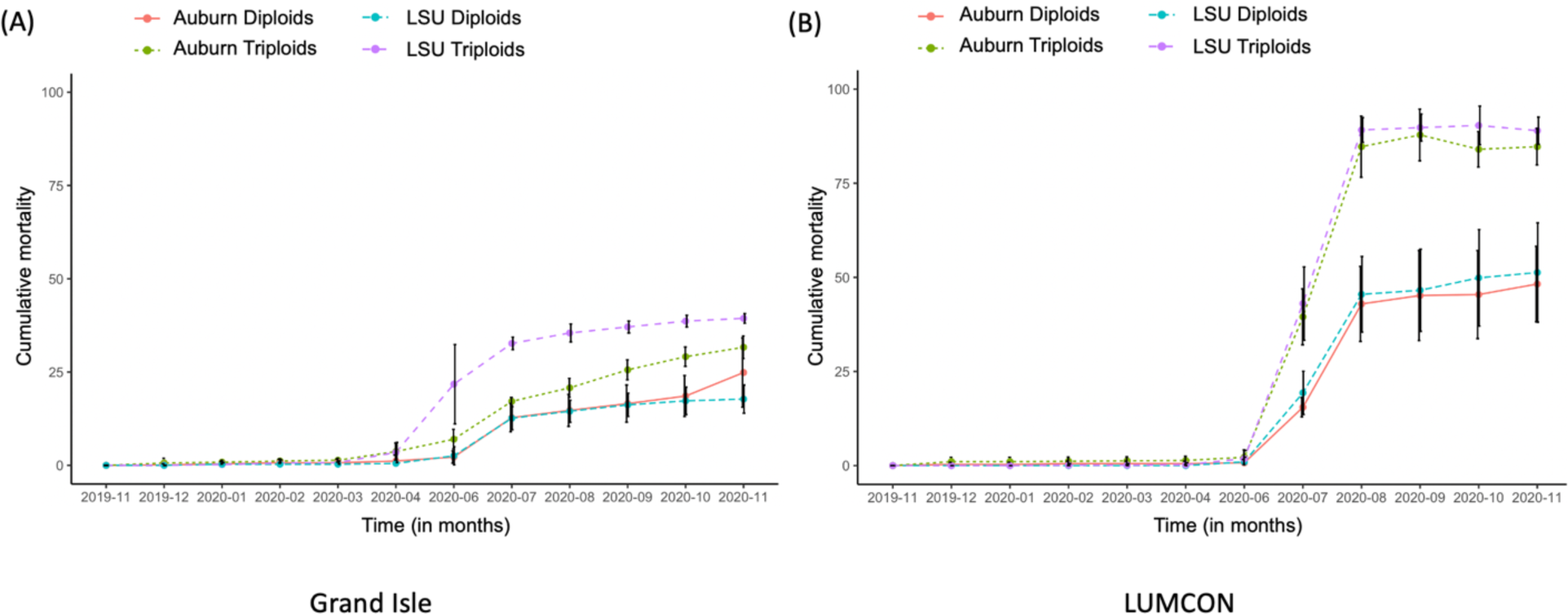
Cumulative mortality of diploid and triploid oysters outplanted to (A) Grand Isle and (B) LUMCON sites, replotted from Bodenstein et al., 2023. The X-axis indicates timepoints in months for which mortality data was collected, and the Y-axis represents cumulative mortality. Oysters are grouped together based on their ploidy and cohorts. Purple line: triploid oysters from LSU cohort, green dotted line: triploid oysters from Auburn cohort, blue dotted line: diploid oysters from LSU cohort, orange line: diploid oysters from Auburn cohort.

**Figure 3:**
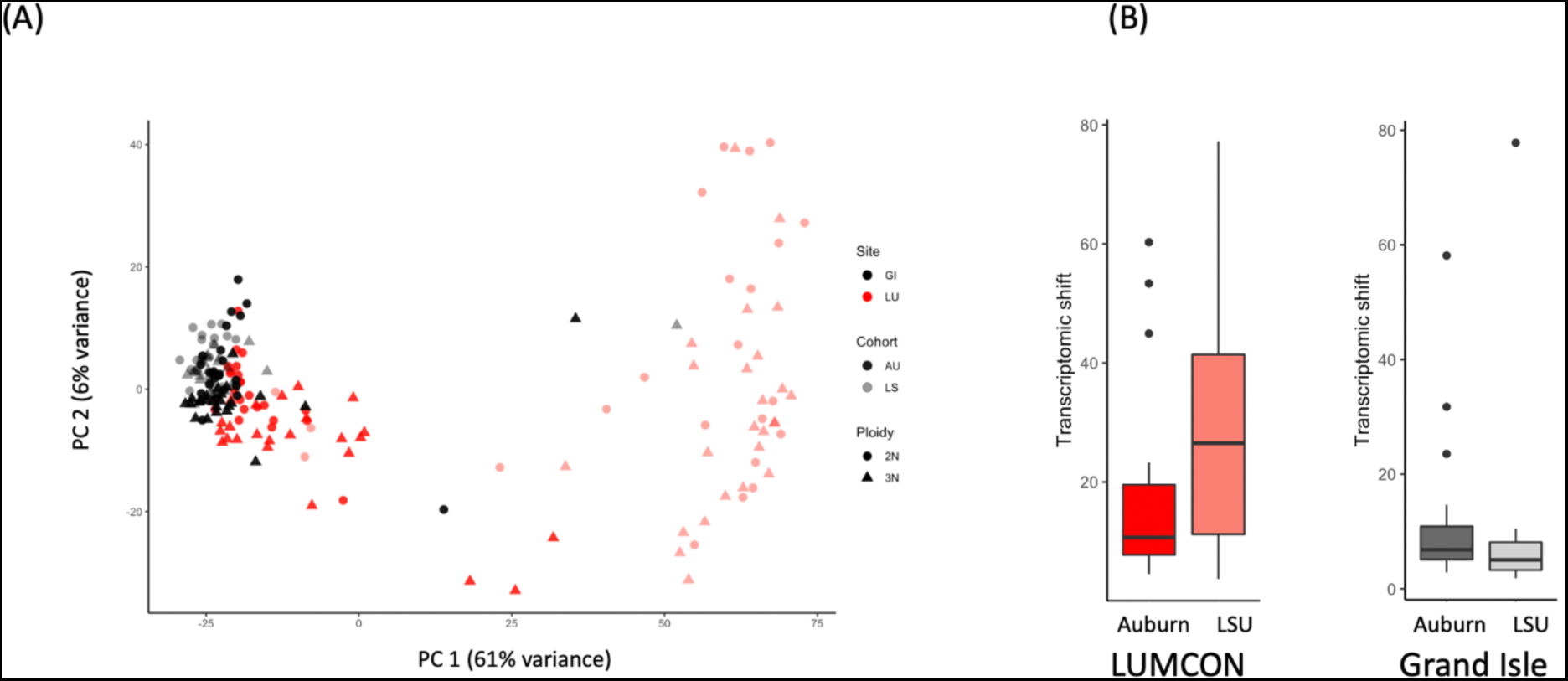
A) PCA showing separation based on site, cohort, and ploidy; B) transcriptomic shift measured as distance in multivariate space between diploid and triploid samples matched for their site, cohort, and dams.

### 3.3 Differential Gene Expression analysis

The results of our DEG analyses are summarized in Table 1. Our pairwise comparisons of diploid vs triploid oysters for sites and cohorts detected 1233 DEGs for LUMCON site and 867 DEGs for LSU cohort. Go analysis performed for these DEGs showed enrichment of GO terms related to calcium ion binding and microtubule-based process for the LUMCON site. On the other hand, no GO terms were detected for Grand Isle site, LSU, and AU cohort. Our pairwise comparisons of diploid and triploid oysters at LUMCON (low salinity) site, for LSU and Auburn cohorts revealed higher number of DEGs for LSU cohort (1812) than Auburn cohort (709).

**Table 1:** Number of DEGs detected for each pairwise comparison performed using EdgeR, DESeq2, and NOISeq. R package VennDiagram was used to find common DEGs detected by all three methods for each comparison.

| Comparison | EdgeR | DESeq2 | NOISeq | Common DEGs | Up-regulated DEGs | Down-regulated DEGs |
| --- | --- | --- | --- | --- | --- | --- |
| 2n LSU vs 3n LSU at GI | 817 | 1860 | 702 | 289 | 113 | 176 |
| 2n LSU vs 3n LSU at LUMCON | 4683 | 5522 | 2864 | 1812 | 1000 | 812 |
| 2n AU vs 3n AU at GI | 689 | 1925 | 198 | 125 | 26 | 99 |
| 2n AU vs 3n AU at LUMCON | 1886 | 2652 | 1628 | 709 | 273 | 436 |
| 2n AU vs 2n LSU at LUMCON | 10690 | 10180 | 6246 | 3779 | 503 | 3346 |
| 2n AU vs 2n LSU at GI | 1368 | 2232 | 1274 | 785 | 612 | 173 |
| 3n AU vs 3n LSU at LUMCON | 10476 | 11810 | 13621 | 9022 | 5437 | 3585 |
| 3n AU vs 3n LSU at GI | 1594 | 1371 | 2657 | 1065 | 677 | 388 |
| 2n vs 3n at GI | 787 | 1371 | 776 | 281 | 117 | 164 |
| 2n vs 3n at LU | 2589 | 3783 | 2867 | 1233 | 594 | 639 |
| 2n vs 3n at AU | 1352 | 2187 | 2678 | 826 | 368 | 458 |
| 2n vs 3n at LSU | 2282 | 3348 | 2243 | 867 | 310 | 557 |

Triploid oysters from LSU cohort downregulated transcripts coding for genes related to calcium signaling (Calmodulin-a, Histidine-rich calcium binding protein (HRC), fibrillin1 & 2, annexin A4, cadherin-23, and cadherin EGF LAG seven-pass G-type receptor 1(CELSR1)) and cell cycle processes (KIF2A, KIF3C, NEK4, NEK6, CDK1, TNKS1, HAUS augmin-like complex subunit 3, MCM7, CHTF18, SMCs, RTEL1, and MAPKKK). GO analysis for this comparison revealed enrichment of GO terms related to microtubules, cell cycle, and calcium binding in triploid oysters. Our pairwise comparisons of diploid and triploid oysters at GI (high salinity) site, for LSU and Auburn cohorts also revealed higher number of DEGs for LSU cohort (286) than Auburn cohort (125). GO analysis performed on LSU + GI DEGs revealed enrichment of GO terms related to reactive oxygen species and superoxide metabolic process by triploid oysters.

GO analysis for Auburn + LUMCON and Auburn + GI DEGs yielded no GO terms. When compared within their ploidies, triploid oysters showed a higher number of DEGs across sites and cohorts than diploid oysters. Across two outplant sites, triploid oysters showed downregulation of transcripts that were enriched for GO terms related to cell cycle regulation and amino acid metabolism and upregulation of transcripts that were enriched for GO terms related to ion transport. Across two cohorts, triploid oysters showed upregulation of transcripts that were enriched for GO terms related to apoptosis and downregulation of transcripts that were enriched for GO terms related to DNA replication.

### 3.4 WGCNA

We also performed Weighted Gene Correlated Network Analysis (WGCNA) to identify groups of genes displaying similar expression patterns in response to site, ploidy, and cohort. Fig. 4a shows 11 modules labeled as color names and the degree of their correlation with traits of interest. We found five (grey60, brown, red, and blue) modules associated with site and two (midnightblue and blue) with cohort. The module blue showed significant association with both site and cohort, while module cyan was associated with site, ploidy, and cohort. Fig. 4b shows eigengene expression plots for brown, grey60, red, cyan, greenyellow, and blue modules with lines indicating samples grouped by their ploidies and cohorts. Of interest are the expression patterns in cyan module, where eigengene expression was higher for triploid oysters, but oysters from Auburn cohort overall showed higher eigengene expression regardless of their ploidies.

**Figure 4:**
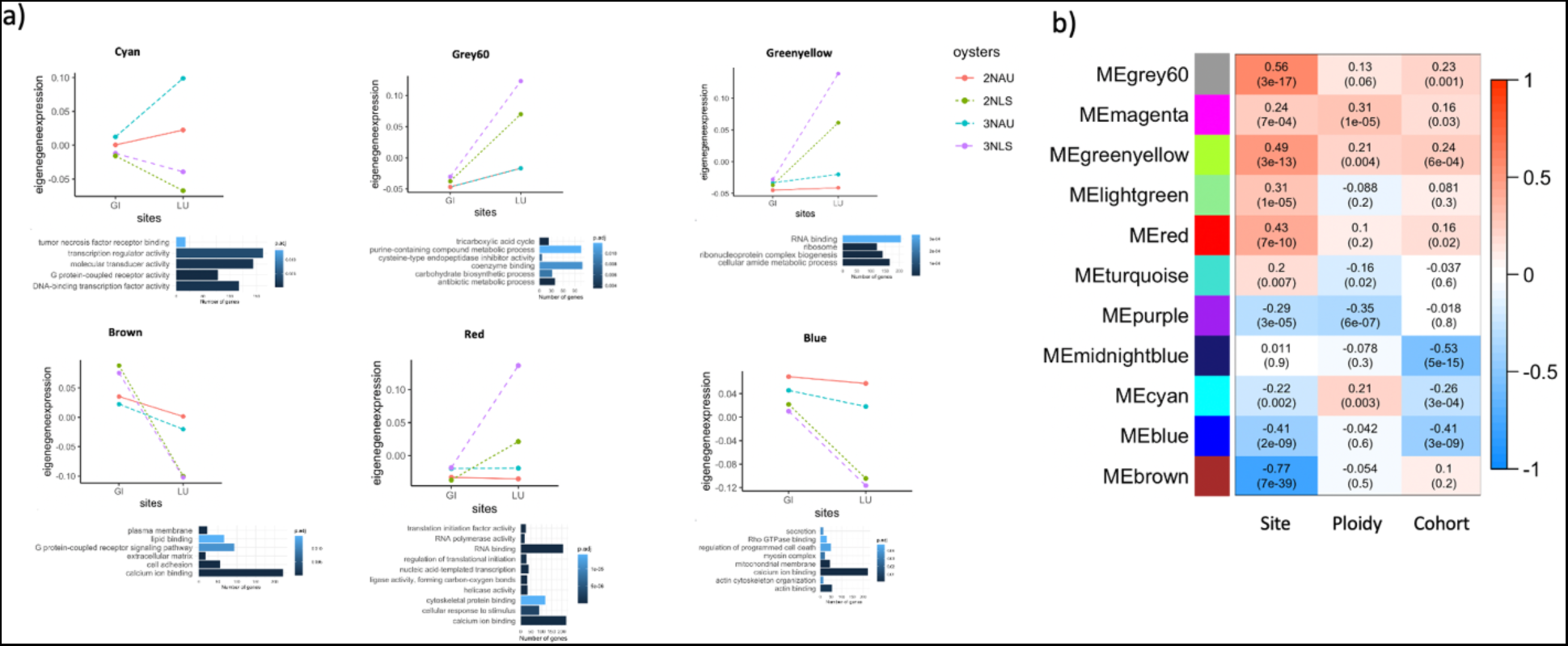
a) Eigengene expression plots for four modules showing significant correlation to site and cohort in WGCNA, with samples grouped together by their cohorts and ploidies; and GO terms with highest p-values in those corresponding modules. Orange: diploids from Auburn cohort, green: diploids from LSU cohort, blue: triploids from Auburn cohort, purple: triploids from LSU cohort. b) Heatmap of WGCNA module-trait relationship results. The X-axis shows the experimental parameters of interest used for correlations. Red indicates positive correlations and blue indicates negative correlations between a module and experimental parameter.

Similarly, eigengene expression in the red module was higher in oysters belonging to LSU cohort but remained unchanged for oysters from Auburn cohort. Red module was positively correlated to site and GO terms overrepresented in this module were related to initiation and regulation of translation, calcium ion binding, cytoskeleton protein binding, and cellular response to stimulus. The blue module was negatively correlated with site and cohort and contained genes that were downregulated by oysters from LUMCON site and LSU cohort. GO analysis for this module revealed an overrepresentation of GO terms related to calcium ion binding and actin binding. The brown module also showed strong negative correlation for site, i.e. genes from this module were downregulated by oysters from LUMCON site when compared with oysters from Grand Isle. GO analysis revealed enrichment of GO terms related to plasma membrane, lipid binding, cell adhesion, calcium binding, and extracellular matrix. On the other hand, Grey60 module was positively correlated to site, and GO analysis for this module revealed overrepresentation of GO terms related to TCA cycle and cysteine type endopeptidase inhibitor activity among others. Patterns seen in blue, brown, and gry60 modules were in agreement with our pairwise DEG comparisons.

## 4. Discussion

The primary aim of our study was to understand molecular and physiological causes of elevated summer mortality in triploid oysters. Contrary to our initial hypothesis, mortality data from our experiment revealed no effect of maternal broodstocks on triploid mortality, indicating that maternal stocks used to breed oysters weren’t the key factor contributing to summer mortality in triploid oysters. Our analyses also did not detect instances of aneuploidy, which has been previously observed in polyploids organisms (Comai, 2005). On the other hand, mortality data showed a strong effect of cohort, which was also reflected in our transcriptomic analysis in the form of DEGs. Our comparisons of diploid and triploid oysters across outplant sites and cohorts revealed a significant number of DEGs and GO terms related to key biological processes, indicating the stronger influence of these two factors on summer mortality. Our inter - and intra - ploidy analyses reveal that diploid and triploid oysters share at least some of their cellular response to low salinity and summer conditions, as seen through enrichment of common GO terms and transcripts. The results from DEG comparisons also match with the correlation patterns observed in WGCNA - the modules significantly associated with site and cohort were enriched for biological processes detected in DEG comparisons.

### Intra-ploidy variation in transcriptomic response to salinity/outplant sites

Our within ploidy comparisons, performed separately for diploid and triploid oysters revealed that cell division was significantly affected in both diploid and triploid oysters at the low salinity site, which experienced greater mortality. We observe that at low salinity site, diploid as well as triploid oysters from the higher mortality LSU cohort upregulate transcripts that were enriched for GO terms related to DNA replication and repair, cell cycle regulation, and response to stimulus. Presence of these common GO terms indicates that regardless of their ploidies, both diploid and triploid oysters respond to low salinity cues by altering or arresting their cell division. Upregulation of genes involved in cell cycle checkpoints in the gill tissue has been previously reported in mussels (Lockwood & Somero, 2011) and killfish (Kültz & Avila, 2001; Whitehead et al., 2011), indicating that cell cycle regulation is a conserved response to low salinity across species. Changes in the salinity have multi-faceted impact on a cell – they can change the osmotic balance inside the cell, activate the cellular signaling pathways that detect changes in salinity, and affect structure and functions of proteins and nucleic acids in the cell. Oysters are osmoconformers, however ability to osmoconform to abrupt and large changes in salinity, however, may take days and is a function of the initial salinity (i.e., salinity oysters are held prior to a change is salinity) and the rate, magnitude and direction of the salinity change (Shumway, 1996; S. C. Hand & Stickle, 1977; M. La Peyre et al., 2003; M. K. La Peyre et al., 2009; McFarland et al., 2013).

Therefore, response to salinity stress has been proposed to be a complex process that targets multiple aims including the initiation of a damage control response to repair the protein and DNA damage caused by salinity change, and also initiating a response to remove molecules or cells that have been damaged beyond repair (Hochachka and Somero, 2002). The enrichment of DNA repair GO terms we see for LSU cohort therefore suggests that they were employing similar stress response mechanisms to combat low salinity stress. Arresting growth during salinity stress is a way of ensuring that damaged DNA isn’t passed down in the cell division cycles – it gives the cell enough time to repair the DNA damage and then progress to cell division. The upregulation of cell cycle checkpoints, which are essentially ‘quality check’ points to ensure the structural integrity on the DNA before progressing to the next step, are also an indicator of LSU triploids coping with low salinity stress.

This common response across ploidies presents an interesting case as in our next set of comparisons, we observe that when compared to each other, triploids up-regulate these transcripts less than diploids. The shared transcriptomic response to salinity and its decreased magnitude in triploids, could mean that triploids mount a weaker adaptive response to salinity, resulting in greater cellular damage. This pattern, where less stress-tolerant genotypes mount a weaker adaptive transcriptomic response to a stressor, is a phenomenon that has been observed in other transcriptomic studies seeking to understand physiological mechanisms underlying intraspecific variation in stress tolerance (DeBiasse & Kelly, 2016)

Along with these shared patterns, we also observed some ploidy-specific gene expression patterns for triploid oysters. At LUMCON, triploid oysters from the LSU cohort upregulated transcripts involved in immune response compared to their Auburn counterparts. In bivalves, abiotic stressors such as temperature and salinity have been shown to induce transcriptional changes related to immune response (Ellis et al., 2011; Gagnaire et al., 2006; Lacoste et al., 2002; Place et al., 2012). The upregulation we observed in the triploid oysters at the low salinity LUMCON could potentially be due to the stressful salinity and temperature conditions they experienced.

### Inter-ploidy variation in transcriptomic response to salinity/outplant sites

Despite finding many DEGs, we did not find any GO terms for oysters from LSU cohort at Grand Isle and for oysters from Auburn cohort at LUMCON. The remaining two categories did yield DEGs as well as GO terms. At Grand Isle, triploid oysters from AUBURN cohort upregulated transcripts enriched for GO terms related to reactive oxygen species metabolic process and superoxide metabolic process.

At LUMCON, triploid oysters from the LSU cohort downregulated genes involved in three key processes - calcium signaling, ciliary activity, and cell cycle checkpoints. This cohort specific response was also seen by Bodenstein et al 2023 in their mortality measurements performed on these oysters – mortality levels for triploids from LSU cohort were highest at low salinity site.

The GO terms and transcripts enriched for these processes hints at some degree of crosstalk among these three processes, suggesting a coordinated stress response in response to salinity, led by calcium signaling. Intracellular calcium is noted for its role as a messenger molecule that facilitates intra and intercellular communication. Specifically, increase in intracellular calcium in response to abiotic stress is a hallmark of plant stress response, and a similar increase in calcium and its receptors (partners) in response to ocean acidification has been noted in the Pacific oyster as well (Xin et al., 2022). Unlike their diploid counterparts, triploid oysters were unable to upregulate genes involved in calcium signaling, indicating triploid-specific problems in detecting and responding to low salinity stress. In our study, triploid oysters from LSU cohort downregulated several key members of calcium-mediated stress response pathway such as Calmodulin-a, Histidine-rich calcium binding protein (HRC), and cadherin EGF LAG seven-pass G-type receptor 1(CELSR1). Calmodulin-a, is a well-studied calcium-binding protein that influences key physiological processes like cell proliferation, differentiation, and apoptosis by interacting with its many downstream targets (Xin et al., 2022). Histidine-rich calcium binding protein (HRC), which is important for calcium homeostasis and cadherin EGF LAG seven-pass G-type receptor 1(CELSR1), a type of G protein-coupled transmembrane receptors instrumental for communication between extracellular and intracellular environment, were also downregulated by triploid oysters (Arvanitis et al., 2011; X.-J. Wang et al., 2014).

Furthermore, genes encoding calcium dependent proteins responsible for structural elements of intra and extracellular space like fibrillin1 & 2, annexin A4, cadherin-23 were also downregulated by triploid oysters. Changes in cell shape and volume do occur in response to salinity, and downregulation of these cytoskeleton elements by triploid oysters may have hindered their effort to maintain their cell shape and osmotic balance under low salinity conditions (Zhang et al., 2017), Hochachka and Somero, 2002). Taken together, downregulation of these calcium related genes shows that low salinity had a domino-like effect on triploid oysters, affecting several key biological processes in them.

Numerous genes coding for cytoplasmic and axonemal dyneins were also downregulated by triploid oysters. As one of the three major cytoskeletal motor proteins, cytoplasmic and axonemal dyneins have been shown to be associated with sensory and motility functions of the cilia, respectively. Upregulation of transcripts involved in ciliary activity at low salinity has been observed in diploid *C. virginica*, and has been hypothesized to help in maintaining homeostasis and avoiding valve closure (Jones et al., 2019). The lack of upregulation of these genes in triploid oysters, combined with their increased mortality at the low salinity site hints at perturbation of ciliary activity in their gill tissue, and future experiments measuring ciliary activity in triploid oysters under summer stress would be helpful to understand the role it plays in triploid mortality. Additionally, morphological measurements performed on these oysters by Bodenstein et al reveal that triploids have larger gill area than diploids, after correcting for their size differences (Bodenstein et al., 2023). Bigger cell size and cell volume of triploid cells has been hypothesized to cause disruption of ion channels and decrease their sensitivity to environment, and negatively affect volume regulation under fluctuating salinity (Guo & Allen, 1994; Haim-Abadi et al., 2023). It is possible that the difference in gill area is due to triploids having larger cells than diploids and is in turn at least partially responsible for disruption of their cellular homeostasis under low salinity stress. Changes in ciliary activity as a response to salinity stress have been observed in marine bivalves (Johnson et al., 2021; Maynard et al., 2018), although we are unclear as to why we observe this for only one cohort of triploid oysters in our study. It is possible that triploid oysters from LSU cohort were more severely affected by salinity due to differences in their rearing conditions as compared to their Auburn counterparts.

Along with ciliary activity, triploid oysters from LSU cohort also downregulated genes involved in cell cycle and its regulation. Genes such as KIF2A, KIF3C, NEK4, NEK6, CDK1, TNKS1, HAUS augmin-like complex subunit 3, and MAPKKK which play key roles in different stages of cell cycle including but not limited to maintenance of chromosome structure, DNA replication and repair, formation and maintenance of spindle fiber, normal progression of cell division, and cell cycle checkpoints (Chang & Karin, 2001; Fry et al., 2012; Hsiao & Smith, 2008; Kültz & Burg, 1998; Lawo et al., 2009; Sheng et al., 2018) were downregulated by triploids at low salinity site. Genes with regulatory functions during cell division like MCM7, which ensures that the replication of chromosomal DNA occurs only once before cell division and CHTF18, which ensures fidelity of transmitting chromosomes from one generation to next were also downregulated in triploid oysters (Berkowitz et al., 2012; Onesti & MacNeill, 2013). Lastly, genes involved in DNA repair (SMCs and RTEL1) were also downregulated by triploid oysters at low salinity site.

Downregulation of several key players involved in cell cycle indicates that gill tissue cells of triploid oysters were encountering errors at multiple stages of cell division. As mentioned in the section about intra-ploidy comparisons, upregulation of genes involved in cell cycle was a shared transcriptomic response observed in both ploidies, but its magnitude appears to be lower in triploid oysters when compared to diploids. This difference in magnitudes could be due to polyploid nature of triploid oysters. Cell division in polyploid cells, especially those with odd number of chromosomes such as triploids is prone to errors such as chromosome loss and aneuploid cells (Comai, 2005). Additionally, progression through cell cycle does depend on environmental stimuli like thermal and osmotic stress (Hochachka and Somero, 2002). Taken together, it appears that the triploid oysters at low salinity sites were particularly challenged with the dual threat of unfavorable environmental conditions and unfavorable cellular milieu for optimal cell division, and therefore had higher mortality rates than diploids.

The cohort effect in our interploidy comparisons could be attributed to differences in the two cohorts. At the low salinity site, field mortality measurements from Bodenstein et al., 2023 show higher cumulative mortalities for triploids from LSU cohort, but do not see a cohort-specific effect on diploid mortality. The cohort effect was also observed in growth rates – at low salinity sites, both diploids and triploids from LSU cohort had slower growth rates. Since the point of distinction between two cohorts was hatchery conditions, we think that early life experience could be influencing growth and survival in later life. Stressors present in the early life environment of oysters have been shown to have an impact on their later life survival (Donelan et al., 2021; Hettinger et al., 2012; Spencer et al., 2020), and the cohort specific differences in growth rates and survival we see could be resulting due to different rearing conditions experienced by oyster larvae at the two hatcheries.

The elevated mortality of triploids oysters observed in our study was marked by enrichment of genes and GO terms related to several cellular processes such as cell division, calcium signaling, and ciliary activity, all of which are also influenced by changes in external environment. As triploid oysters have been shown to lose their growth advantage over diploids in stressful environments (Callam et al., 2016; Davis, 1994; Wadsworth et al., 2019b) and are also more vulnerable to multi-stressor scenarios (George et al., 2023; J. Brianik & Allam, 2023; Li et al., 2022), direct physiological measurements that track these processes during the summer conditions, would be helpful for determining the exact causes of the triploid mortality.

Additionally, as our sampling time was during peak summer (July 2020), both diploid and triploid oysters in our study were exposed to environmental spawning cues. It is possible that triploid oysters in our study responded to spawning cues by investing in gametogenesis, and the addition of this metabolically costly process while experiencing abiotic stress threw their energy budget into a disarray. As we used gill tissue for our transcriptomics analyses, we were unable to detect gene expression changes related to reproduction in our experiment. Future transcriptomic studies using gonadal tissue samples will be helpful to detect transcriptomic signatures of reproductive effort in triploid oysters. Furthermore, triploid oysters in our study did show gonadal development but were not tested for formation of viable gametes. This distinction is important, as previous studies have shown that even though triploid *C. virginica* develops gonads, it fails to produce viable gametes and shows errors in synchrony of gametogenesis (Matt & Allen, 2021). The cost of this unsuccessful reproductive effort and its impact on survival of triploid oysters remains unexplored. Future studies aimed at exploring tradeoffs of this imperfect gametogenesis in triploids will be helpful in clarifying the contribution of reproduction in triploid mortality. Lastly, previous studies have shown occurrences of aneuploidy and subsequent chromosome loss in triploid oysters, both of which can cause cell death by disrupting normal patterns of gene expression (De Sousa et al., 2016). Even though we did not detect any incidents of aneuploidy, our test was an indirect measure and was performed on gill tissues of the diploid and triploid oysters. Future studies involving direct measures like karyotyping gonadal cells of triploid oysters across their developmental stages will be helpful in understanding whether these genomic irregularities exist and contribute to their mortality.

## Acknowledgements

We thank Dr. Brian Callam, F. Scott Rikard, and staff members of the Michael Voisin Oyster Research Lab and Hatchery and Auburn University Shellfish Lab for producing the oysters used in this study.

## Supporting information

Supplementary file 1: Count matrix used in downstream differential gene expression analysis and WGCNA.

Supplementary file 2: Enriched GO terms detected in differential gene expression analysis and WGCNA.

Supplementary table 1: Regression line slopes calculated for ten chromosomes for each diploid and triploid oyster pair, matched for their site, cohort, and dam.

Supplementary Figure 1: PCAs showing separation based on A) Site, B) Cohort, C) Ploidy, D) Dams.

